# Biodiversity mediates the effects of stressors but not nutrients on litter decomposition

**DOI:** 10.1101/2020.03.03.972547

**Authors:** Léa Beaumelle, Frederik De Laender, Nico Eisenhauer

## Abstract

Understanding the consequences of ongoing biodiversity changes for ecosystems is a pressing challenge. Controlled biodiversity-ecosystem function experiments with random biodiversity loss scenarios have demonstrated that more diverse communities usually provide higher levels of ecosystem functioning. However, it is not clear if these results predict the ecosystem consequences of environmental changes that cause non-random alterations in biodiversity and community composition. We synthesized 69 independent studies reporting 660 observations of the impacts of two pervasive drivers of global change (chemical stressors and nutrient enrichment) on animal and microbial decomposer diversity and litter decomposition. Using meta-analysis and structural equation modelling, we show that declines in decomposer diversity and abundance explain reduced litter decomposition in response to stressors but not to nutrients. While chemical stressors generally reduced biodiversity and ecosystem functioning, detrimental effects of nutrients occurred only at high levels of nutrient inputs. Thus, more intense environmental change does not always result in stronger responses, illustrating the complexity of ecosystem consequences of biodiversity change. Overall, these findings provide strong empirical evidence for significant real-world biodiversity-ecosystem functioning relationships when human activities decrease biodiversity. This highlights that the consequences of biodiversity change for ecosystems are nontrivial and depend on the kind of environmental change.

## Introduction

Human activities cause global environmental changes with important consequences for biodiversity and the functioning of ecosystems. Understanding these consequences is crucial for better policy and conservation strategies, which will ultimately promote human well-being too^1^. A key question is to what extent changes in ecosystem functioning are mediated by changes in biodiversity. Extensive research has demonstrated that biodiversity is needed for the stable provenance and enhancement of ecosystem processes and functions^2–4^. However, this body of evidence is mostly based on experiments comparing ecosystem functioning in artificial communities with varying number of species. Such experiments might not capture the complex ways by which shifts in biodiversity induced by global change ultimately affect ecosystem functioning^5,6^.

Early biodiversity-ecosystem function (BEF) experiments typically controlled for environmental gradients, thus not accounting for the underlying drivers of biodiversity change^5,7,8^. These early experiments also focused on species richness as the sole biodiversity index, and manipulated it directly and randomly. However, environmental change will often elicit non-random changes in several facets of biodiversity^9–11^ (community composition and population densities^12,13^, functional diversity^14–16^, trophic diversity^17,18^, or intra-specific diversity^19^). The selective effects of environmental change emerge because different organisms differ in their response to environmental change. For example, larger organisms and predators are often more negatively affected than smaller organisms at lower trophic levels^8,20,21^. In addition, several variables that are not directly related to biodiversity control ecosystem functions (e.g. physiological rates^22^ and alterations of physical and chemical conditions^5,10^). When environmental change affects these mechanisms, teasing out the relative importance of biodiversity-mediated effects is complicated even more. Given the number of different potential mechanisms, quantifying the extent to which shifts in biodiversity underpin the effect of environmental change on ecosystem functioning under real-world scenarios of global change is a key challenge for ecology^5–8,11,23^. Incorporating the impacts of environmental change drivers into BEF studies and meta-analyses is an important step forward to address such questions^5,6^.

The vast majority of BEF experiments has focused on plant richness and ecosystem functions such as biomass production^11^. Litter decomposition has tremendous importance for biogeochemical cycles. A recent study highlighted that wood decomposition releases almost as much CO_2_ to the atmosphere as fossil-fuel combustion^24^. Small changes in the rate of this process could thus have important consequences for the overall carbon balance and climate warming. The diversity of decomposers (invertebrates and micro-organisms that fragment and decompose organic matter in both aquatic and terrestrial systems) is crucial litter decomposition^25–29^ and for ofther ecosystem functions as well^4,30,31^. Despite the importance of decomposers, BEF experiments focusing on litter decomposition more often addressed the influence of plant litter diversity than of decomposers^26,32^. In a meta-analysis, decomposer diversity had greater effect on decomposition than the diversity of plant litter^33^, although also weak and neutral effects have been reported^11^. Facilitation and complementarity through niche partitioning are primary mechanisms underlying the positive relationship between decomposer diversity and decomposition^25,26,32^. The need to quantify environmental change effects on decomposer diversity, along with potential knock-on effects on litter decomposition, is therefore particularly pressing.

There is a variety of environmental change drivers, and different types of drivers may have diverse effects on biodiversity and ecosystem functions^5^. We postulate that there are two main categories of environmental change: stressors (e.g., temperature, drought, chemicals) and resource enrichment (e.g., of CO_2_ or mineral nutrients). While stressors cannot be consumed, and act as conditions that alter growth rates, resources are by definition consumed, which has important implications for how they should enter theory^34,35^. Chemical stressors and nutrient enrichment are important case studies of environmental stressors and resource enrichment, because their presence is increasing rapidly^36^ and they are projected to have severe effects on biodiversity^37^. They are also of particular relevance for decomposer communities. Chemical stressors such as metals and pesticides decrease the diversity, abundance, growth and activity of decomposers across terrestrial and aquatic systems^38–40^. Contrastingly, nutrient enrichment can have positive impacts on the abundance and physiological rates of decomposer organisms by reducing resource limitations^41^, but at the same time decrease decomposer diversity^42,43^. Across ecosystems, stressors and nutrients can exert opposite impacts on litter decomposition rates, with decreases in response to chemical stressors but increases following nutrient enrichment^44,45^. In addition, decomposition involves both microorganisms and invertebrates^25,26,46^ that may respond differently to stressors and nutrients with a higher sensitivity of invertebrates than microorganisms^47,48^. Although many published case studies report shifts in decomposer diversity and in rates of litter decomposition at sites impacted by stressors and nutrients, biodiversity-mediated effects have not yet been quantified across systems.

Here we addressed the question if the effects of stressors and nutrient enrichment on decomposer diversity and abundance explain the response of litter decomposition to these two types of pervasive environmental change drivers (Fig. 1). We synthesized 69 published case studies reporting the impact of stressors (metals, pesticides) and nutrients (nitrogen or phosphorous additions) on litter decomposition and on decomposer diversity (taxa richness, Shannon diversity, evenness) or abundance (density, biomass) at sites differing in stressor or nutrient levels. Our comprehensive global dataset of 660 observations encompasses studies across taxonomic groups (animal and microbial decomposers), ecosystems (aquatic and terrestrial) and study types (experimental and observational) (Fig. 2). We quantified the effect size of environmental change on decomposer diversity or abundance and on litter decomposition within studies using correlation coefficients between stressor or nutrient levels and decomposer diversity, abundance and litter decomposition. We also characterized stressor and nutrient intensities, and standardized their levels in water, soil or sediment using environmental quality criteria issued by environmental authorities (e.g. ECHA, USEPA, UKTAG). Using meta-analysis and structural equation modelling (SEM), we first compared the overall effects of stressors and nutrients on decomposers and decomposition across systems and studies (first meta-analysis), and second, addressed to what extent changes in decomposer diversity and abundance mediate the impacts of these two contrasting drivers of environmental change on decomposition (second meta-analysis and SEM). Third, we explored the effects of three main moderators on decomposers diversity, abundance and decomposition responses, as found in the second meta-analysis: stressor or nutrient intensity, taxonomic group (animal vs. microbes) and study type (experimental vs. observational studies).

**Figure 1:**
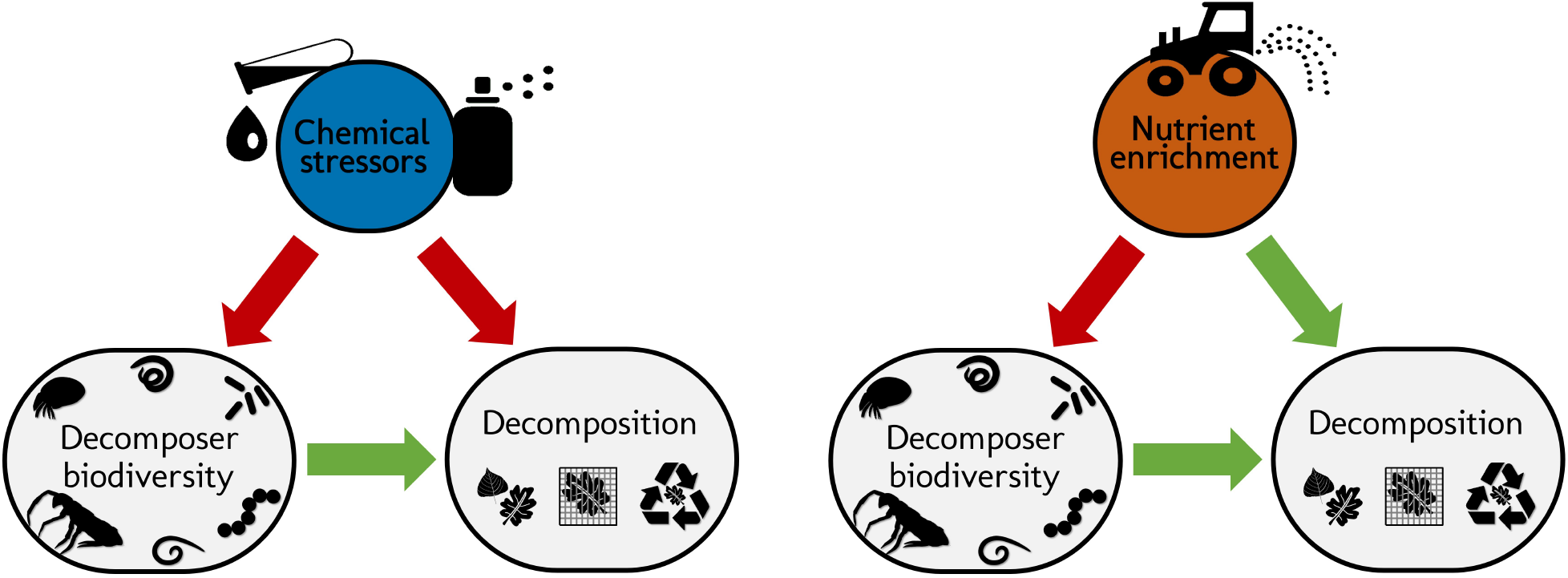
Schematic representation of the structural hypotheses tested in this study. Green arrows depict expected positive effects, red arrows represent negative effects Stressors and nutrients are hypothesized to decrease decomposer diversity. The response of decomposers diversity to environmental change drivers determines the response of decomposition following biodiversity-ecosystem functioning theory (Srivastava et al. 2009). Stressors and nutrients can affect litter decomposition independent of changes in decomposer diversity, especially through changes in physiological activity (De Laender et al. 2016, Giling et al. 2019).

**Figure 2:**
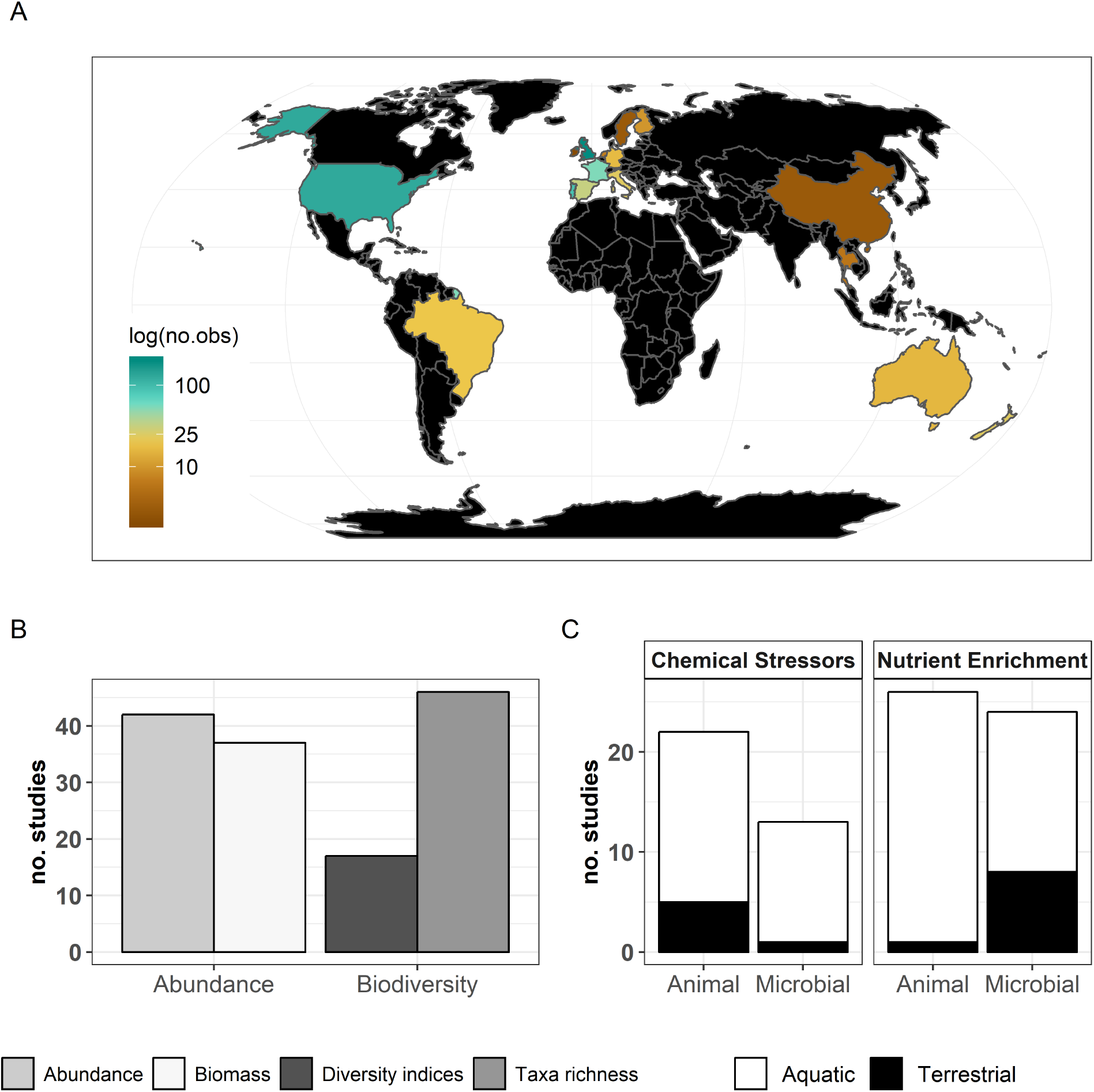
Description of the data used in the present meta-analysis. A: countries represented and corresponding number of observations, B: decomposer diversity and abundance metrics covered, and C: decomposer taxonomic groups and ecosystem types represented.

We expected that chemical stressors and nutrients would have contrasting effects on decomposer diversity and abundance, and on litter decomposition across ecosystems and studies (Fig. 1). We hypothesized that chemical stressors generally decrease decomposer diversity, abundance^40,49^ and litter decomposition rates^45,48^, and that nutrients generally decrease decomposer diversity^42,43^ but increase decomposer abundance and litter decomposition rates (based on physiological effects and decreasing resource limitations^41,42,44^). We further hypothesized that litter decomposition responses to environmental change depends on changes in decomposer diversity and abundance, and expected an overall positive relationship independent of environmental change intensity based on BEF theory^33^.

## Results

### Description of the data and overall patterns

The final dataset contained 69 (case) studies from 59 publications, representing 660 observations. Data were mostly from Europe (44; 443 (studies; observations)) and North and South America (19; 168), while Asia (2; 9) and Oceania (4, 40) were less represented (Fig. 2A). The studies covered aquatic (55; 388) and terrestrial systems (14; 272) (Fig. 2C), and used observational (43; 336) or experimental approaches (26; 324). Studies reported abundance (66; 463) or diversity responses (48; 197) (Fig. 2B) of soil and benthic invertebrates (48; 509) and microbes associated with litter materials (36; 151) (Fig. 2C). Chemical stressors were mostly metals (13; 257) and pesticides (12; 66) associated with industrial activities, accidental spills, and agricultural practices. Nutrient enrichment studies addressed fertilization by various N and/or P forms (26; 175), and eutrophication due to agricultural runoffs (10; 59) or wastewater effluents (4; 44). There was no study reporting nutrient enrichment impact on soil decomposer diversity in the dataset. Funnel plots and intercepts of Egger’s regression showed evidence for positive publication bias in nutrient enrichment studies reporting decomposer abundance (Fig. S2, Table S2). No publication bias was detected in the other datasets.

We found largely contrasting effects of stressors and nutrients on each of the three response variables, in a first-level meta-analysis comparing the overall effects of the two drivers of environmental change (Fig. 3, Table S3). Chemical stressors overall decreased decomposer diversity, abundance and litter decomposition across studies (Fig. 3). Nutrient enrichment tended to decrease decomposer diversity but to increase abundance and decomposition, although these trends were not significant as indicated by confidence intervals of the grand mean effects overlapping with zero (Fig. 3).

**Figure 3:**
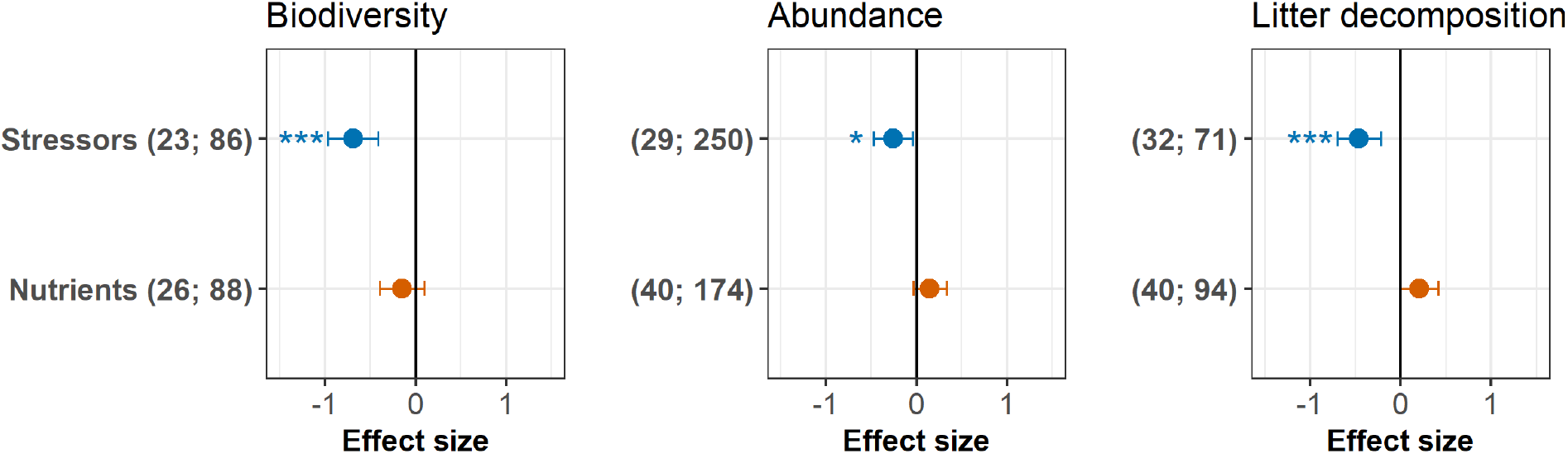
Grand mean effect sizes of chemical stressors and nutrient enrichment on decomposer diversity (taxa richness and diversity indices), abundance (density and biomass), and litter decomposition. Effect sizes are z-transformed correlation coefficients. Error bars show 95% confidence intervals. Numbers in parentheses indicate number of studies and observations, respectively. Symbols show the significance level (∗ ∗ ∗ p < 0.001; ∗ p < 0.05). For full model results, see Table S3.

### Biodiversity-mediated effects of stressors and nutrients on litter decomposition

The responses of decomposition and of decomposer diversity and abundance to chemical stressors were correlated: decreases in decomposition were associated with decreases in decomposer diversity and abundance (Fig. 4 upper panels). We did not find such a relationship for nutrients. Instead, a range of positive and negative responses of decomposer diversity, abundance, and decomposition was found, without significant association between them (Fig. 4 lower panels). In addition, there was a wide range of decomposition responses when decomposer diversity and abundance responses were close to zero (intercepts from Fig. 4 lower panels).

**Figure 4:**
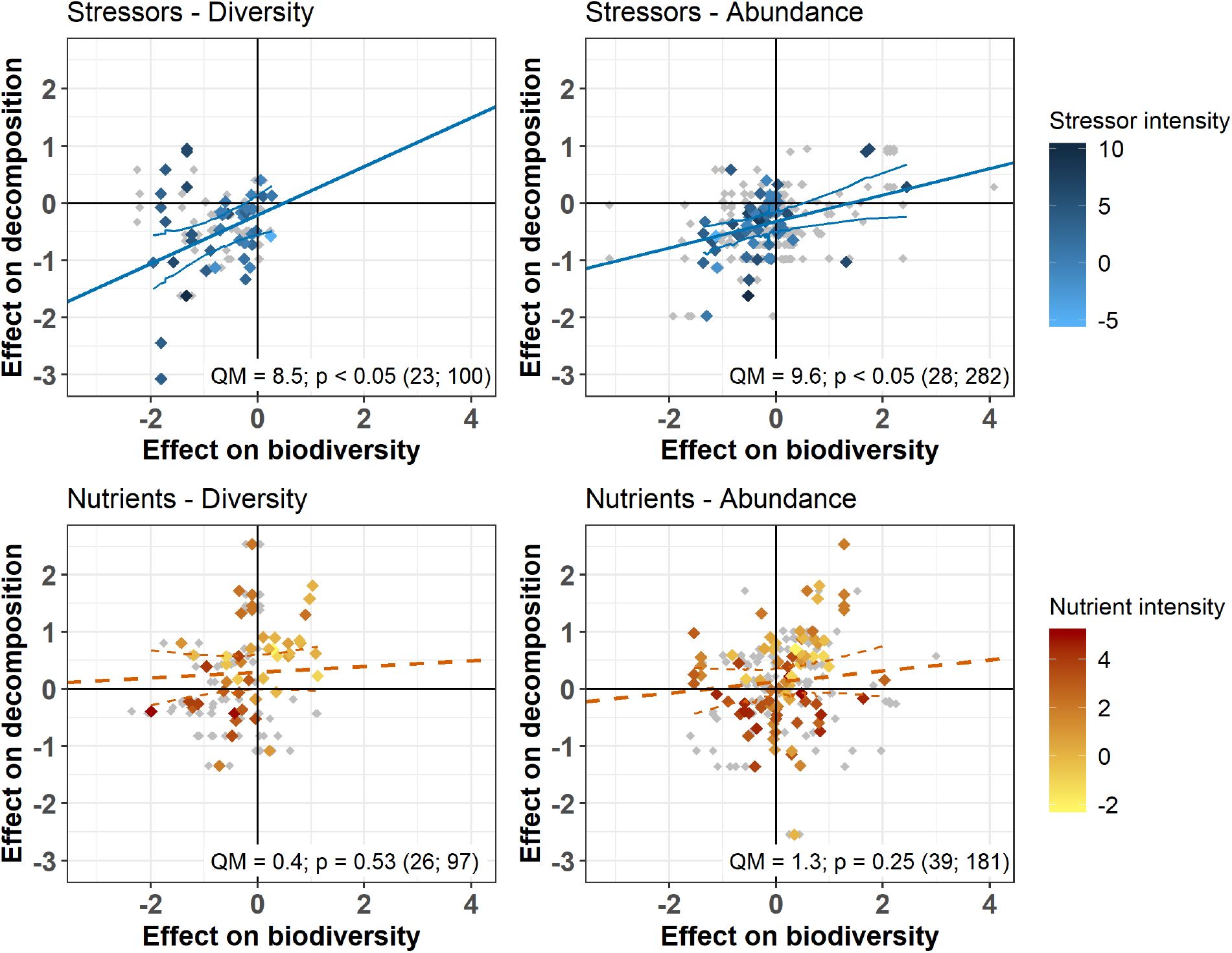
Relationship between the responses of decomposition and decomposer diversity and abundance to chemical stressors and nutrient enrichment. Variables are effect sizes (z-transformed correlation coefficients) of stressors or nutrients on litter decomposition and on animal and microbial decomposer diversity (left panels) or abundance/biomass (right panels). Gray symbols are individual observations of effect sizes; Colored symbols indicate the mean effect size on biodiversity or abundance across effect sizes on litter decomposition. Darker colors represent a higher standardized level of environmental change. Lines represent meta-regressions between effect sizes for decomposition and decomposers, where solid lines are statistically significant (p < 0.05), dashed lines are non-significant (p > 0.05), and thin lines depict the regression’s confidence interval. Panels show the statistics of the meta-regressions with sample size (number of studies; number of observations).

According to our overarching hypothesis, the SEM indicated that the effects of stressors on litter decomposition were mediated by shifts in decomposer diversity and abundance. Including the direct paths from decomposer diversity or abundance to litter decomposition improved both the models according to mediation tests and AIC comparisons (Fig. 5, Table S4). In addition, the path coefficients from diversity and abundance to the decomposition response to stressors had (standardized) values higher than 0.1 (Fig. 5) and were statistically different from zero (Table S5). However in contrast to chemical stressors, the SEM did not support biodiversity-mediated effects of nutrient enrichment on litter decomposition. While the mediation test and AIC indicated that the decomposer diversity-mediated path improved the model (Fig. 5, Table S4), the path coefficient was not significantly different from 0 (Table S5). The decomposer abundance-mediated path of nutrients was not supported by the data: an SEM without the direct path from decomposer abundance to decomposition could not be rejected based on the mediation test (Fig. 5, Table S4), and including this path did not improve the model according to the AIC comparison. Besides, we found publication bias in this dataset (Fig. S2, Table S2), and model check indicated that the residuals of the nutrients-abundance model were non-independent from the fitted values. The results from this model are reported here for comparison purposes only.

**Figure 5:**
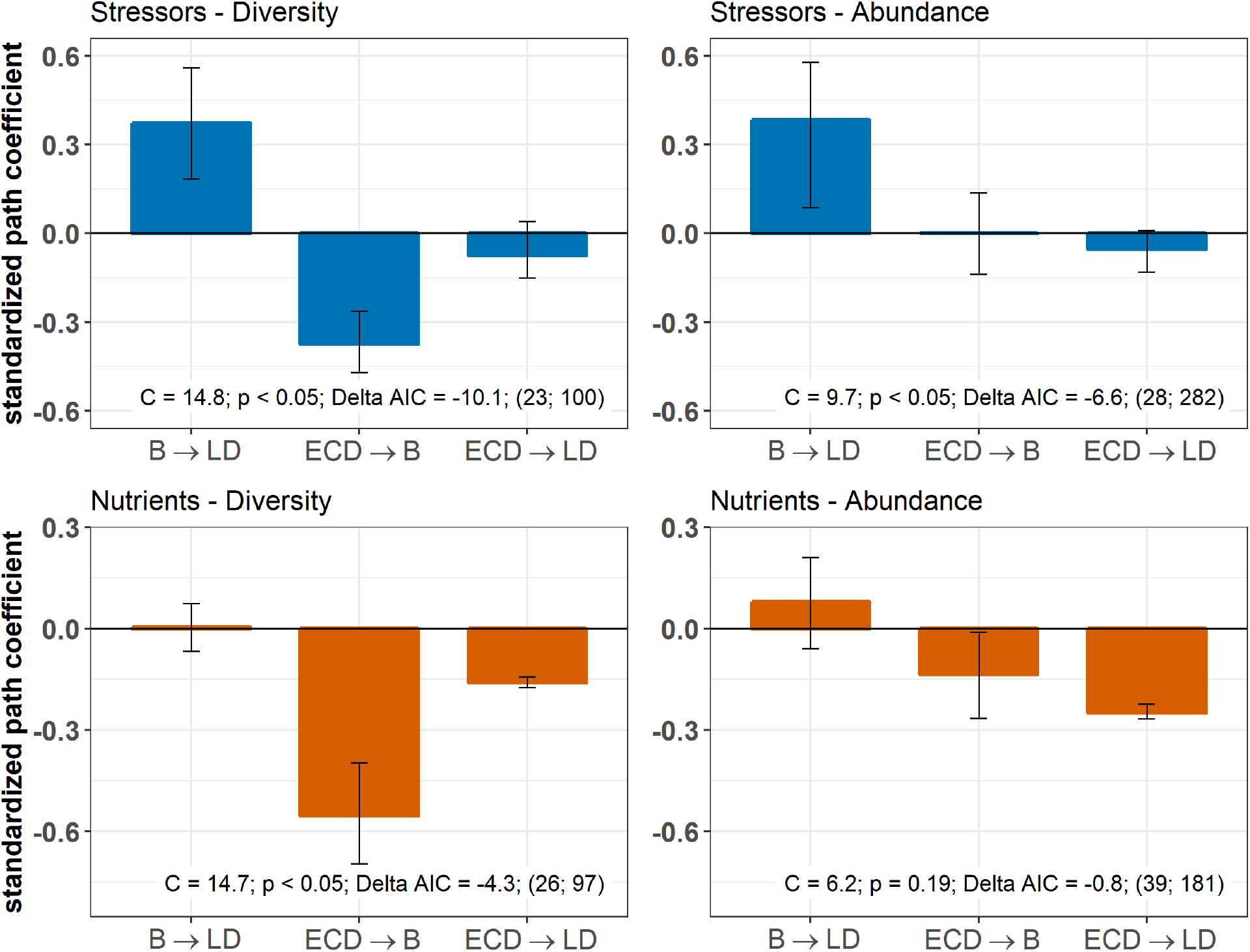
Decomposer diversity and abundance explained litter decomposition response to chemical stressors but not to nutrient enrichment. Biodiversity-mediated effects of environmental change drivers (chemical stressors and nutrient enrichment) on litter decomposition (B LD) and direct effects on decomposition (ECD LD) and decomposer diversity or abundance (ECD B). Values are standardized path coefficients from structural equation models relating effect sizes of stressors and nutrients on decomposition and on decomposer diversity in terms of taxa richness, Shannon or evenness (left panels) or decomposer abundance and biomass (right panels). Bars and error bars are the means and 95% confidence intervals of model coefficients derived with data resampled randomly (see Table S5 for unstandardized path coefficients). Results of mediation tests: C statistics and p-values are significant (p < 0.05) when models omitting the path B LD were not consistent with the data. Delta AIC is the difference in AIC score between models with and without B LD path. Sample sizes are reported (number of studies; number of observations).

The magnitude of the biodiversity-mediated effects of chemical stressors on decomposition was stronger than that of the direct effects of stressor intensity on decomposition. The total indirect effect of stressors on decomposition mediated by diversity (i.e. the mathematical product of standardized paths B *→* LD and ECD *→* LD; Fig. 5, Table S6) was twice that of the direct effect of stressors on decomposition (indirect diversity-mediated path = - 0.14, direct path = - 0.07, Table S5) while the abundance-mediated effects was negligible (indirect abundance-mediated path = 0.0003, direct path = - 0.05, Table S6). In the case of nutrient enrichment, however, decomposition responses were not explained by shifts in decomposer diversity and abundance, and the direct effects of nutrient intensity dominated the total effect (indirect diversity-mediated path = - 0.003; direct path = - 0.16; indirect abundance-mediated path = - 0.01; direct path = - 0.25, Table S6). Finally, between model comparisons based on the unstandardized path coefficients^50^ revealed that decomposer diversity was a stronger driver of decomposition response to stressors than decomposer abundance (B *→* LD path of 0.42 and 0.24 respectively for diversity and abundance, Table S6).

Sensitivity analyses revealed that the results were robust to the inclusion of approximated standard deviations (Fig. S3, Tables S8, S9), and extreme values of effect sizes (Fig. S4, Tables S10, S11). We found partially different results when using log response ratios as effect sizes (Fig. S5, Tables S12, S13), due to lower sample sizes and emergence of extreme values in these datasets. In addition, the log-response ratio is probably sensitive to the various metrics of biodiversity, abundance, and decomposition covered by the individual studies that we included, while correlation coefficients better accomodate such discrepancies^51^.

### Response of animal and microbial decomposers and decomposition to stressor and nutrient intensity

Despite the overall negative effects of stressors on decomposition, negative responses in decomposition were not associated with higher stressor intensity (Fig. 5, 6). This result held for two complementary approaches: multivariate SEM (Fig. 5) that relied on data resampling to account for replicated values of decomposition matching several decomposer responses (e.g. for different taxa in the same litterbag), and meta-regressions (Fig. 6) where data resampling was not necessary (see Methods). Decomposer diversity responses decreased with stressor intensity according to the SEM (Fig. 5) but this trend was not significant in the bivariate meta-regression (Fig. 6). Decomposer abundance responses were not associated to stressor intensity in both the SEM and meta-regression approaches (Fig. 5, 6). We found different patterns for nutrient enrichment, where decomposition responses decreased with nutrient intensity (Fig. 5, 6), from positive effects at low intensity to negative effects at higher intensity (Fig. 6). A similar pattern was observed for decomposer diversity where responses decreased with nutrient intensity from positive to neutral to negative responses at high nutrient levels (Fig. 6). Nutrient intensity however did not explain the responses of decomposer abundance (Fig. 5, 6) and both positive and negative responses were found at high nutrient levels.

**Figure 6:**
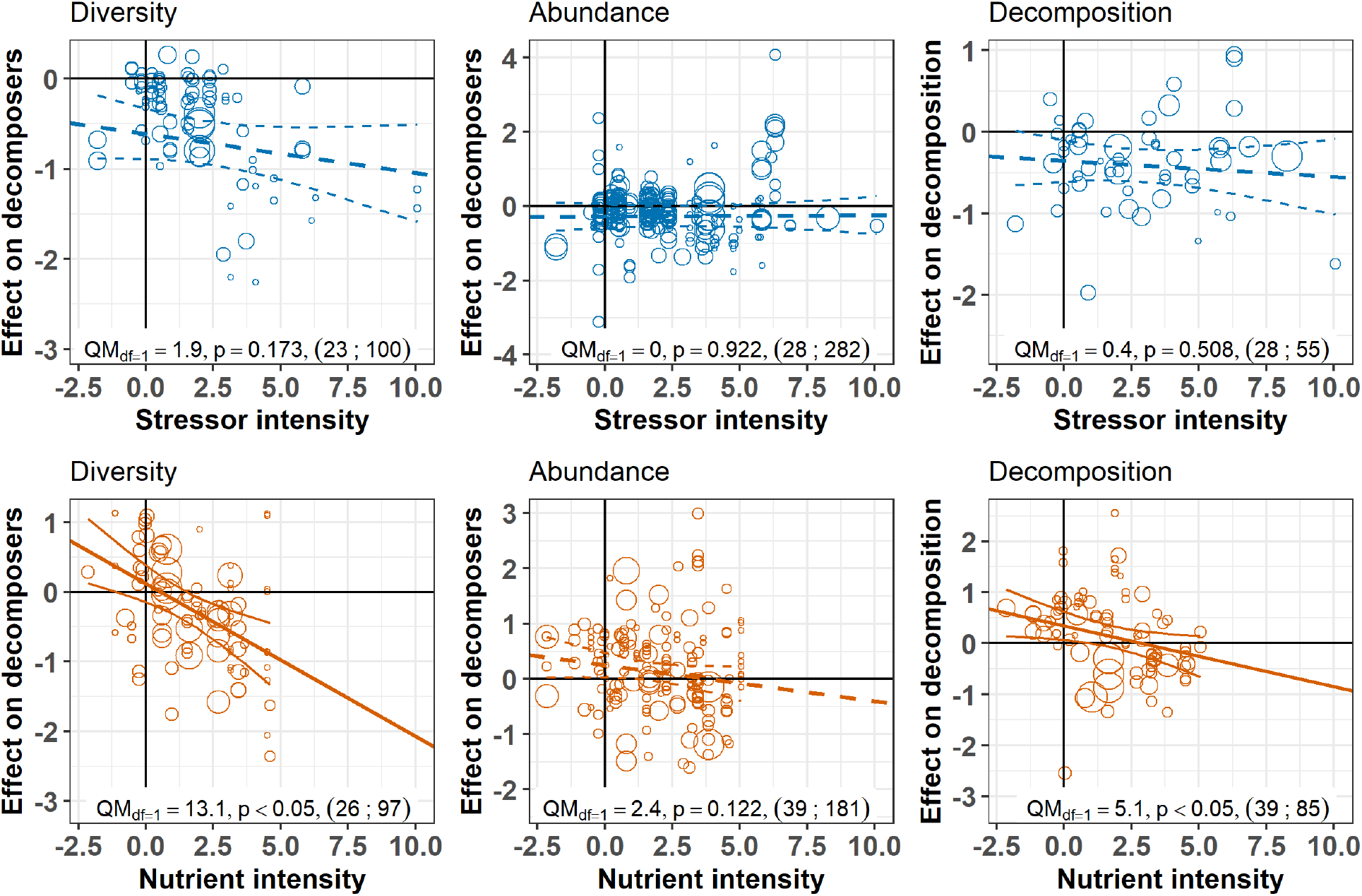
Decomposer and decomposition responses to the intensity levels of chemical stressors and nutrient enrichment. Values are effect sizes (z-transformed correlation coefficients). Stressor or nutrient intensity represents the standardized level of environmental change in the treatment with the highest level (values < 0: observed level below quality criteria considered to be safe for the environment; values > 0: observed level above quality criteria). Point size is proportional to the inverse of the variance in effect size. Lines are the slopes and 95% confidence intervals from bivariate meta-regressions, with associated QM statistics, p-value and sample size (number of studies; number of observations).

The meta-analysis further revealed clear discrepancies between the response of animal and microbial decomposers to stressors and nutrients. Animal decomposers responded more strongly to chemical stressors than microbial decomposers. The mean effects of chemical stressors on animal decomposer diversity and abundance were more negative than that on microbial decomposers, confirmed by Wald type tests of the second-level meta-analyses (Fig. 7 upper panels, Table S7). Animal decomposers overall decreased in diversity but increased in abundance in response to nutrient enrichment (Fig. 7, lower panels). On the other hand, the mean effects of nutrients on microbial decomposer diversity and abundance had lower magnitude compared to animals (Table S7), with confidence intervals overlapping with zero (Fig. 7 lower left panel). Finally, there was no clear difference between observational and experimental studies (Fig. 7, Table S7), and between biodiversity responses in terms of taxa richness or of diversity indices (Table S7).

**Figure 7:**
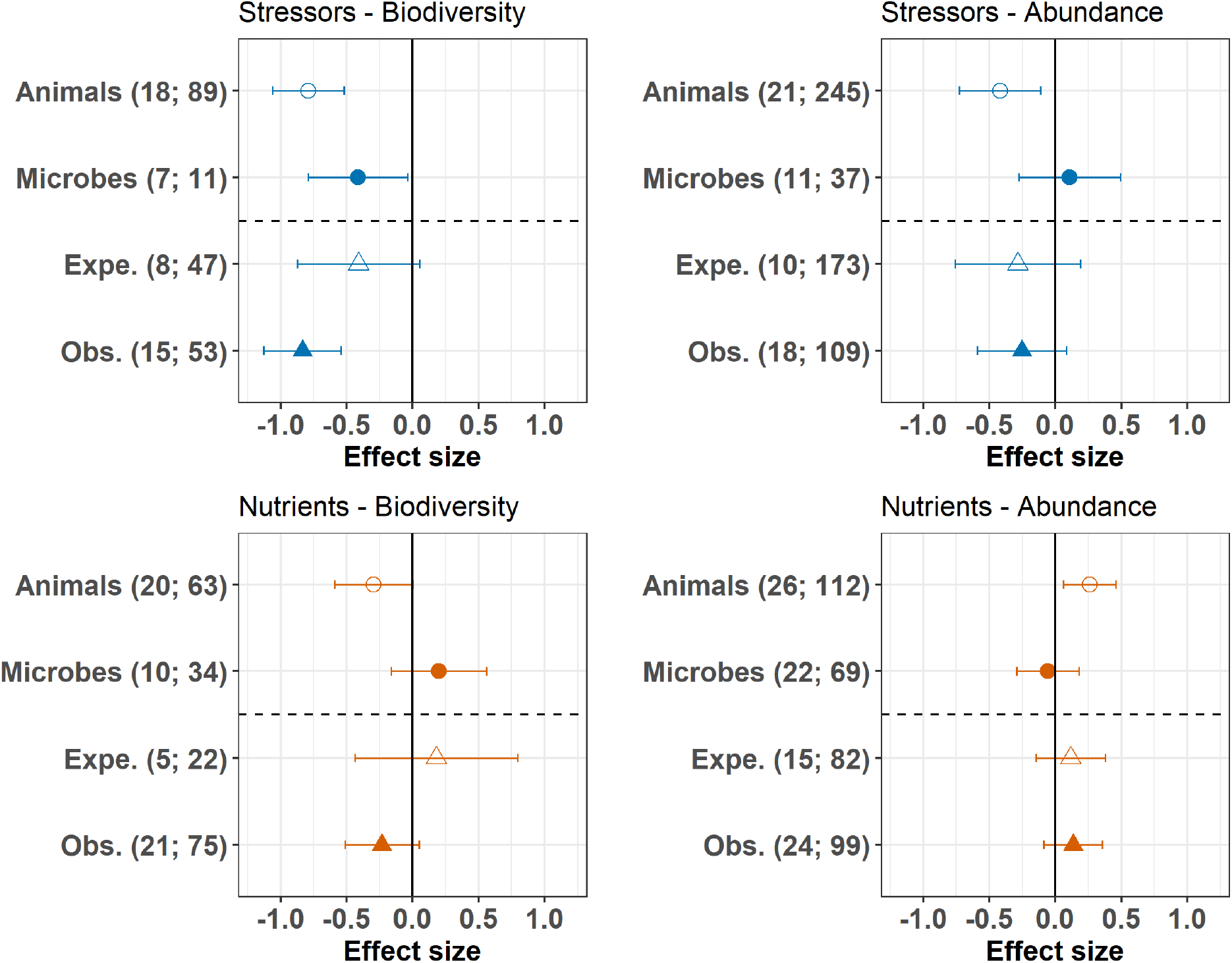
Moderator effects on decomposer diversity and abundance and on decomposition responses to chemical stressors and nutrient enrichment. Responses of decomposer diversity (taxa richness and diversity indices) and abundance (densities and biomass) to stressors and nutrients according to the taxonomic group (animals and microbes) and study type (Expe. = experimental; Obs. = observational studies). Values are mean effect sizes (z-transformed correlation coefficients) and 95% confidence intervals derived from meta-analytic models. Sample sizes are reported for each moderator: (number of studies; number of observations).

## Discussion

The present synthesis brings new insights into how changes in decomposer biodiversity induced by two pervasive drivers of environmental change ultimately affect decomposition. We find concomitant changes in biodiversity and decomposition under the influence of chemical stressors but not nutrient enrichment, highlighting that real-world patterns relating shifts in biodiversity and ecosystem functioning depend on the type of environmental change. In fact, we observed significant correlations between effects on biodiversity and ecosystem function in a scenario where chemical stressors caused significant decline in biodiversity. By contrast, in cases where nutrient enrichment caused variable responses in biodiversity, relationships between biodiversity and ecosystem function responses were weaker. It remains an understudied but important question if results of controlled BEF experiments are applicable to non-random changes in biodiversity caused by human activities^5–8,11,23^. The present results provide strong empirical evidence for significant real-world BEF relationships when human activities decrease biodiversity.

### Biodiversity-mediated effects of chemical stressors on decomposition

Chemical stressors caused consistent reductions in decomposer diversity and abundance as well as in litter decomposition rates, in line with several previous case studies^52,53^ and meta-analyses^45,48^. Adding to the previous knowledge, the present meta-analysis shows that changes in decomposer diversity and abundance explained the decomposition response to stressors, providing evidence for the expectation that shifts in biodiversity mediate the impact of chemical stressors on decomposition. We acknowledge that despite the SEM analysis, the approach conducted here remains correlative. However, our study builds on a body of experimental and observational evidence that already demonstrated that more diverse and abundant decomposer communities support higher decomposition rates, albeit not under the influence of environmental change^27,28^.

We especially complement a previous meta-analysis showing the importance of decomposer diversity for decomposition across experiments manipulating the richness of invertebrate and microbial decomposer communities^33^. We extend on this and show that non-random biodiversity loss induced by stressors are closely associated with decreases in decomposition across a wide range of studies. A recent review pointed out that in naturally-assembled terrestrial communities, studies more often found neutral and to a lesser extent positive relationships between decomposer diversity and decomposition^11^. In that review, communities were not influenced by environmental change drivers, and the vote counting approach used is sensitive to the statistical power of individual studies and could have increased the probability of finding non-significant relationships^51^. In line with our findings, an experiment mimicking the sequence in which freshwater invertebrate decomposers are lost after disturbances showed that decreasing non-randomly the number of species decreased decomposition rates^54^.

Biodiversity-ecosystem function experiments manipulating biodiversity directly are key to understand the mechanisms involved in this relationship^9^, especially because they control for the effects of environmental heterogeneity or abundance. However, in real-world scenarios, environmental change drivers affect both biodiversity and abundance simultaneously. As demonstrated here, this is especially the case for stressors impacts that decrease decomposer diversity and abundance^40^. The abundance or biomass of different decomposers is of critical importance for decomposition^55–57^. Even at constant richness and community composition, strong decreases in abundance can have important impacts on ecosystem functioning^12,58^. It is beyond the scope of the present meta-analysis to disentangle the effects of biodiversity from the effects of abundance, and we found that both contributed to explain shifts in decomposition in separate analyses. It is interesting to note that the few cases where negative effect sizes of stressors on biodiversity were associated with positive effect sizes on decomposition were also cases where decomposer abundance was positively associated with stressors (Fig. 4). Although we cannot specifically test this with the present data, it seems that in those particular cases, increases in decomposer abundance counteracted the negative effects of decreases in decomposer diversity^59^. These concomitant shifts in both diversity and abundance have important implications for our estimates of diversity responses, as studies mostly reported richness to estimate decomposer diversity, but rarely corrected for the sampling effort^60^. This means that lower abundances rather than a lower number of species *per se* might have directly caused some of the negative effects on biodiversity reported here^61^. This common caveat in meta-analysis approaches that rely on how individual studies report biodiversity, also applies to the present study, and reinforces the importance of reporting rarefied richness in future studies of the impacts of chemical stressors on biodiversity and ecosystem functioning.

The effects of changes in decomposer diversity and abundance on decomposition found in the present study might also have channeled changes in community and food-web structure not captured by our biodiversity metrics. Changes in keystone species^25^, functional diversity^14,15^, vertical diversity^17,26^, or dominance patterns^62^ might have shifted concomitantly to taxonomic diversity and abundance. Unfortunately, studies rarely reported these together with decomposition measurements. For example in our dataset, only 7 studies reported evenness. Future studies need to explore shifts in decomposer community composition in more detail to better understand what particular aspect of biodiversity is responsible for changes in decomposition rates^10,25^. For example, there is ample evidence that shifts in functional diversity are crucial for decomposition^14^, and that facilitative interactions occur primarily between decomposers of contrasting body size^32^. This is especially the case for interactions between animal and microbial decomposers, where fragmentation of litter by detritivores facilitates access for microbial decomposers^25,63^

Here, we found that invertebrates were more affected by chemical stressors than microbes, across aquatic and terrestrial ecosystems. Invertebrate decomposers are particularly sensitive to the impacts of metals and pesticides^38–40,48^. This is consistent with the general expectation that larger organisms are more sensitive to environmental change due to longer generation time, higher energetic demands and lower population densities^21,64,65^. These different sensitivities between groups of decomposers could imply that the biodiversity-mediated effects of stressors on decomposition are more strongly linked to shifts in invertebrates than microbes such as reported in a previous review^48^. However, in another meta-analysis focusing on microbial-driven decomposition rates, changes in fungal biomass and richness explained shifts in decomposition under the impacts of chemical stressors, but also of nutrient enrichment^43^.

### Nutrient-induced changes in decomposition were not related to shifts in decomposer diversity

Nutrient enrichment had contrasting impacts on litter decomposition and decomposer diversity compared to chemical stressors confirming our expectations. These different biodiversity and function responses led to different emergent relationships between decomposer diversity and decomposition compared to stressors. We found that nutrients had a variety of effects ranging from positive to negative depending on the taxonomic group (Fig. 7) and nutrient intensity (Fig. 6), and resulting in neutral overall mean effects (Fig. 3). Previous syntheses also found positive^44^ as well as inconsistent^66^ responses of decomposition rates to nutrient enrichment in streams. The relatively small mean effect of nutrient enrichment on decomposition in the present meta-analysis could be explained by the use of correlation as an effect size, which does not capture potentially non-monotonic responses of decomposition to nutrients^42^. However, we noted that most of the studies included in the present meta-analysis did not individually span nutrient gradients sufficiently large to capture this potential non-monotonous response. Taken together, the studies show positive effects on decomposition at low nutrient intensities that shifted towards neutral to negative effects at higher intensities (Fig. 6), which is consistent with previous findings^42,44^ Low nutrient intensities might have enhanced microbial activity and biomass by alleviating resource limitation, resulting in enhanced decomposition. At higher intensities, however, negative impacts on invertebrate decomposers might have decreased decomposition rates^42,48^.

Contrary to our expectation, nutrient-induced shifts in decomposer diversity and abundance were not associated with shifts in decomposition rates across studies. We found that increasing nutrient intensity decreased the effects on decomposition and on decomposer diversity, but not on decomposer abundance. Statistically controlling for the joint effect of nutrient intensity with SEM indicated no residual association between shifts in decomposer diversity or abundance and in decomposition rates, i.e. a non-significant BEF relationship. Changes in microbial abundance in response to nitrogen deposition explained the responses of different ecosystem functions in terrestrial systems in previous meta-analyses^41,67^. Here we show that this pattern cannot be generalized across aquatic and terrestrial systems and animal and microbial decomposers. Contrary to stressors, when decomposer diversity and abundance were not affected by nutrients, we observed large positive and negative shifts in decomposition (intercepts of Fig. 4), that were explained by nutrient intensity (Fig. 4: negative effects on decomposition at invariant biodiversity are associated with high intensities and positive effects with lower intensities). Together, these results show that nutrient-induced shifts in decomposer diversity were not as strong drivers of decomposition changes as stressor-induced biodiversity shifts. These differences may be partly due to the different mechanisms underlying the effects of stressors and nutrients. Based on previous studies, we speculate that our results are due to the complex responses of animal and microbial decomposers at different nutrient intensities.

Animal decomposers showed a stronger response to nutrients than microbes. Invertebrate decomposers overall decreased in diversity, but they increased in abundance under nutrient enrichment. These results could reflect a loss of sensitive taxa to the benefit of tolerant taxa that were able to use additional resources and would then increase in density. Overall, microbial decomposers responded little to nutrient enrichment probably reflecting a mixture of positive and negative effects that nutrients can have on microbial growth^41,43^, as well as on different microbial taxa. Indeed, nutrients can alleviate resource limitations at low intensities, but can also exert toxic effects at high intensities. The initial levels of nutrients thus condition subsequent responses in decomposers and decomposition to nutrient enrichment^44,66^. Furthermore, at high intensities, nutrients can be associated with other chemical stressors (e.g. pesticides in agricultural runoffs)^42,44^. The influence of interactive effects of stressors and nutrients was impossible to quantify with the data at hand given that only a few experiments assessed the effects of both drivers independently, but many observational studies may have been confounded by such joint effects. Chemical stressors and nutrients are often co-occurring in e.g. agricultural landscapes, and the consequences of such combinations are still poorly understood. Furthermore, stressor and nutrient effects might be modulated by climatic and other environmental conditions, and studies on interaction effects are scarce^22^.

In conclusion, this study brings new insights into the real-world patterns relating ecosystem function to non-random changes in biodiversity induced by environmental change. We found that the consequences of changes in biodiversity for ecosystem functioning depend on the type of environmental change. Real-world scenarios do not necessarily involve concomitant changes in both biodiversity and function across terrestrial and aquatic systems. We further found that at the environmental quality criteria used in risk assessment, there were already significant positive and negative effects on decomposers and decomposition (intercepts of Fig. 6), highlighting the need to better incorporate biodiversity and ecosystem function into ecological risk assessment programs^68^. Finally, we report overall negative effects of chemical stressors on biodiversity and ecosystem functioning across terrestrial and aquatic ecosystems that reinforce recent calls to consider chemical stressors as important global change drivers and address their impacts on biodiversity and ecosystems^36,37^. Positive real-world BEF relationships may be particularly significant in cases where environmental changes decrease biodiversity, such as in the case of chemical stressors. Such information are crucial if we are to design policy and conservation strategies able to reconcile human development with biodiversity conservation.

## Material and Methods

### Data collection

We searched the Web of Science for studies that addressed the impact of environmental drivers and recorded decomposer community responses and litter decomposition rates. The search strategy is fully reported in Supplementary Methods. The search retrieved 2536 references. Abstracts and titles were screened to identify a final set of 61 records that met our inclusion criteria (PRISMA plot, Fig. S1). To be included in the meta-analysis, studies had to:

- Report litter decomposition (rates, mass loss, proportion of mass remaining) and the diversity, abundance, or biomass of decomposers at sites differing in chemical stressor or nutrient levels.
- Focus on naturally-assembled communities subjected to the impact of chemical stressors or nutrient enrichment. Studies that manipulated decomposer diversity directly were not considered to only focus on non-random biodiversity change scenarios. We included mesocosm studies only when they used field-sampled communities and left time for the community to reach an equilibrium in mesocosms in order to reflect real-world conditions as much as possible.
- Report the response of animal (benthic macroinvertebrates, or soil micro, meso or macrofauna) or microbial decomposers (bacteria or fungi from decomposing leaves or in surrounding water or soil samples).
- Report decomposer abundance (density or biomass), or decomposer diversity (taxa richness, Shannon diversity, evenness).

When a reference reported different environmental change drivers or geographical areas with a specific reference site for each case, we considered these as individual (case) studies^67^. We extracted means or sums, standard deviations and sample sizes of litter decomposition, decomposer diversity and abundance (outcomes) in non-impacted vs. impacted sites (control-treatment studies), or at each site when gradients of chemical stressors or nutrients were investigated (gradient studies). When response variables were reported at different time points, we kept only the last time point to capture long-term responses. For studies reporting decomposition, decomposer abundance or diversity for several litter types (e.g. different litter species), several groups of organisms (e.g. functional feeding groups for macroinvertebrates), and several diversity metrics (e.g. Shannon indices and taxon richness), we created separate observations within case studies. We also extracted chemical stressor or nutrient levels at those sites (water, soil or sediment concentrations of chemical stressors or nutrients, or application rate of pesticides or fertilizers). The study type (experimental vs. observational), taxonomic group (animal decomposers or microbial decomposers) and metric of diversity (taxa richness or diversity indices (Shannon diversity and evenness)) were also recorded. We used the online software Webplotdigitizer to extract data from figures^69^. We converted standard errors and confidence intervals into standard deviations using the equations in 70. When reported as mass loss, litter decomposition data were transformed into k rates using the exponential decay equation used in 44.

### Effect size calculation

We used z-transformed correlation coefficients as effect sizes in order to cope with the heterogeneity of data and study types^51^. For control-treatment studies, we first calculated Hedge’s d, and then transformed Hedge’s d into correlation coefficients^70^. For gradient studies (4 or more treatment levels), we calculated correlation coefficients between the mean values of abundance, diversity, or decomposition rate and the corresponding chemical stressor or nutrient concentrations. When means, standard deviations, or sample sizes were missing, we contacted the authors to retrieve the data. When the information could not be retrieved, standard deviations were approximated from the data, using the linear relationship between mean values and standard deviations across our datasets^70^.

### Standardization of chemical stressors and nutrient enrichment intensities

Given the variability in the different stressors and nutrients combinations in the studies, stressor and nutrient levels were standardized into a common environmental change driver intensity (*ECD*_*intensity*_) as follows:

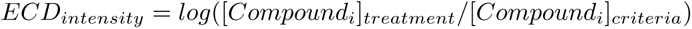

where [*Compound*_*i*_]_*criteria*_ were environmental quality criteria set by European or US environmental authorities for the chemical stressor or nutrient considered (Table S1), and [*Compound*_*i*_]_*treatment*_ were the concentrations of the chemical stressor or nutrient at the treatment or impacted sites. When multiple stressors or nutrients were reported, we used the standardized intensity of the stressor or nutrient corresponding to the highest standardized intensity for the rest of the analyses.

### Overall effects of chemical stressors and nutrient enrichment: first-level meta-analysis

We first tested the differences between the effects of chemical stressors and nutrient enrichment on decomposer diversity, abundance and litter decomposition responses by quantifying the grand mean effect sizes on the three response variables (first level meta-analysis). Three separate meta-analyses were conducted, one for each response variable, and included the type of driver (stressors or nutrients) as a categorical moderator, and a random effect of the case study. We used a weighted meta-analysis giving more weight to effect sizes derived from studies with larger sample sizes. Weights were the inverse of the variance in z-transformed correlation coefficients^71^. Publication bias was evaluated using funnel plots with environmental change driver type as covariate. The intercepts from Egger’s regressions (standardized effect size vs. precision = 1/SE) were inspected for significant deviation from zero that would indicate publication bias^51^. Residual plots were used to detect strong deviation from normality and outliers. We estimated the grand mean effect sizes and compared the effect of chemical stressors and of nutrients using Wald-type chi-square tests. The rma.mv() function of the R package metafor was used^71,72^.

### Relationship between biodiversity and decomposition: Structural equation modelling

An SEM was fitted to estimate the relationship between decomposer diversity or abundance and litter decomposition responses to environmental change drivers while controlling for the joint influence of stressor or nutrient intensity and categorical covariates. We used piecewise SEM^73^ estimating two linear mixed effect models, one for decomposition (*z*_*LD*_) and one for decomposer diversity or abundance responses (*z*_*B*_), with a random effect of the case study on the intercepts. These two sub-models embedded in the piecewise SEM were the second-level meta-analyses in our hierarchical approach. The random effect structure, weighting approach and variance structure were coded with the R package nlme^74^ in a way that fully reproduced the meta-analysis approach of weighting and of known residual variance^75^:

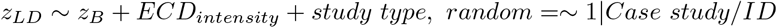

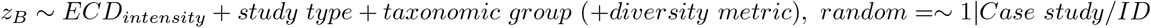

This SEM was tested separately for each of four datasets: Stressors – Biodiversity; Stressors – Abundance; Nutrients – Biodiversity and Nutrients – Abundance datasets. The influence of the diversity metric (diversity indices versus taxa richness) was tested in the Biodiversity datasets only. We initially considered more complicated model structures, but were unable to use them for analysis due to data limitations (in particular the effect of the ecosystem type and of interactions between our covariates).

Outliers, relationships between covariates and non-linear patterns between continuous covariates were explored graphically. Studies often reported different decomposer diversity or abundance values for the same litter decomposition (e.g. when several taxonomic or functional groups were reported in the same litterbag). This variability could have affected the model estimates. We thus used data resampling to account for duplicated effect sizes on litter decomposition in the analyses. A stratified resampling was conducted where for each duplicated value of effect size on decomposition, one randomly selected effect size on biodiversity was kept at each out of 1,000 iterations. The models were fitted for each data resampling iteration and we averaged model estimates and statistics across iterations and used the means as final values (path coefficients and standard error of the path and intercepts, Chi-square statistics and AICs).

Goodness-of-fit of the SEMs was assessed using directed separation tests based on the Fisher’s C statistic. We used mediation tests to explore the significance of the path between decomposer diversity or abundance and litter decomposition based on the Fisher’s C statistic of SEM that did not include the biodiversity-mediated path^73,76^. We calculated the p-value associated with the mean Fisher’s C statistic across data resampling iterations (p-value < 0.05 indicated poor model fit). The AICs of models with and without the biodiversity-mediated paths were further compared using averaged AICs across data resampling iterations. We considered the biodiversity (or abundance) path to be consistent with the data when the SEM without the biodiversity-path had p-value < 0.05 (poor fit) and was not associated with a better AIC value (i.e. lower than 2 units) than the SEM including the biodiversity path. Residuals from the two sub-models of each SEM were graphically evaluated for strong departure to normality and relationship with the fitted values^77^. For these analyses, we averaged the residuals across data resampling iterations for each observation. We finally compared the relative magnitude of the biodiversity-mediated path versus the direct path from stressor or nutrient intensity to litter decomposition based on the mathematical product of the standardized path coefficients^50^.

### Moderator analyses: Second-level meta-analyses

In order to quantify the influence of the categorical (study type, taxonomic group and diversity metrics) and continuous (environmental change intensity) moderators on the three response variables, we further analyzed the results of the second-level meta-analyses (i.e. the sub-models embedded in the SEMs). The data resampling used in the SEM was no longer necessary because there were no repeated values of decomposition matching different decomposer diversity or abundance measurements in this univariate approach. We quantified the effects of the different moderators based on the Wald-type chi-square tests derived with the R package metafor^71^.

### Sensitivity analyses

We finally tested the robustness of the results to the approximation of standard deviations, t he presence of extreme values, and the metric of effect size used. The analyses were re-run with datasets that did not include the effect sizes for which we approximated standard deviations, for datasets that did not include extreme values of effect sizes (values beyond the whiskers of boxplots i.e. below quantile 1 minus 1.5 times the interquartile range or above quantile 3 plus 1.5 times the interquartile range). Finally, we calculated log-response ratios instead of correlation coefficients as effect sizes and re-run the analyses.

## Acknowledgements

L.B. was funded by the Synthesis Centre (sDiv) of the German Centre for Integrative Biodiversity Research (iDiv) Halle-Jena-Leipzig, funded by the German Research Foundation (FZT 118). We are grateful to Helen R. Phillips, Benjamin Rosenbaum, Adam T. Clark and Katharina Gerstner for statistical advice and to Simone Cesarz for creating the images for Fig. 1.

## Authors contribution

All authors conceived the project; L.B. collected the data, performed the analyses and wrote the manuscript; all authors discussed the results and contributed to the manuscript text.

